# Photoimmunotherapy retains its anti-tumor efficacy with increasing stromal content in heterotypic pancreatic cancer spheroids

**DOI:** 10.1101/2021.11.09.467929

**Authors:** Mohammad A. Saad, Wonho Zhung, Margaret Elizabeth Stanley, Sydney Formica, Stacey Grimaldo-Garcia, Girgis Obaid, Tayyaba Hasan

## Abstract

Pancreatic ductal adenocarcinoma (PDAC) is an aggressive disease characterized by increased levels of desmoplasia that contributes to reduced drug delivery and poor treatment outcomes. In PDAC, the stromal content can account for up to 90% of the total tumor volume. The complex interplay between stromal components, including pancreatic cancer associated fibroblasts (PCAFs), and PDAC cells in the tumor microenvironment (TME) have a significant impact on prognoses and thus needs to be recapitulated in vitro when evaluating various treatment strategies. This study is a systematic evaluation of photodynamic therapy (PDT) in 3D heterotypic coculture models of PDAC with varying ratios of patient derived PCAFs that simulate heterogenous PDAC tumors with increasing stromal content. The efficacy of antibody-targeted PDT (photoimmunotherapy; PIT) using cetuximab photoimmunoconjugates (PICs) of benzoporphyrin derivative (BPD) is contrasted with that of liposomal BPD (Visudyne®), which is currently in PDT clinical trials for PDAC. We demonstrate that both Visudyne®-PDT and PIT were effective in heterotypic PDAC 3D spheroids with a low stromal content. However, as the stromal content increases above 50% in the 3D spheroids, the efficacy of Visudyne®-PDT is reduced by up to 10-fold, while PIT retains its efficacy. PIT was found to be 10-fold, 19-fold and 14-fold more phototoxic in spheroids with 50%, 75% and 90% PCAFs, respectively, as compared to Visudyne®-PDT. This marked difference in efficacy is attributed to the ability of PICs to penetrate and distribute within spheroids with a higher stromal content, whereas Visudyne® is restricted to the spheroid periphery. This study thus demonstrates how the stromal content in PDAC spheroids directly impacts their responsiveness to PDT and proposes PIT to be a highly suited treatment option for desmoplastic tumors with particularly high degrees of stromal content.

## Introduction

Pancreatic ductal adenocarcinoma (PDAC) has dismal survival rates (9%), the lowest amongst all cancer types^1,2^. It is refractory to most therapies, including chemotherapy and immunotherapy^3-5^. One hallmark of pancreatic ductal adenocarcinoma (PDAC) is the desmoplastic response that results in a dense, fibrous stroma in the tumor microenvironment (TME)^6-8^. This process is facilitated by activated cancer associated fibroblasts (CAFs). The CAFs and the associated dense stroma contribute to the hypovascular nature of PDACs, reduced drug delivery and distribution, and resistance to therapies thus leading to poor treatment outcomes^8,9^. Tumor stroma is typically comprised of extracellular matrix (ECM) components, CAFs, immune cells, and other tissue-specific cells. The complex interactions between cancer cells and the stroma in the TME play a critical role in tumor growth and metastasis^6,7^. Therefore, there is a significant need for *in vitro* tumor models that mimic the stroma observed in PDAC to design and evaluate therapies for overcoming this barrier.

Several studies have demonstrated the generation of heterotypic *in vitro* models, consisting of PDAC and other stromal cells, recapitulating the complex cellular heterogeneity observed in these tumors^10^. While such models have been useful in assessing response to therapy, most of these studies utilize fixed ratios of these cell types which is often not the case in PDAC patients, where stromal components have been shown to range from 10 – 90% of the total tumor volume^11-17^. In the context of this present study, Tanaka et al. have reported the generation and characterization of an *in vitro* PDAC spheroid model with tunable proportion of stromal components^18^, which recapitulates the PDAC pathophysiology. Motivated by these models, we developed a heterotypic PDAC model, comprising of PDAC cells (MIA PaCa-2) and Pancreatic Cancer Associated Fibroblasts (PCAFs) with varying ratios and expressing mCherry and EGFP to track cellular responses, for comparing the efficacy of photoimmunotherapy (PIT) and non-targeted Visudyne®-Photodynamic therapy (PDT).

Although stromal components are known to confer resistance to most therapeutic strategies employed for PDAC management, PDT has been shown to overcome this resistance by perturbation of stromal and cellular components thereby resensitizing tumor cells to subsequent therapies^19-23^. PDT relies upon the photochemical properties of a photosensitizer (PS), a nontoxic molecule that can absorb light and undergo intersystem crossing to reach an excited triplet state^24^. The longer-lived triplet state of a PS can lead to the formation of cytotoxic reactive molecular species, including singlet oxygen, to induce cell death and TME modulation through various mechanisms^24-26^. Visudyne®, a clinically approved liposomal formulation of benzoporphyrin derivative (BPD) is known to confer cellular phototoxicity by targeting the mitochondria and endoplasmic reticulum^27^. It is being evaluated in Phase II clinical trials for unresectable pancreatic cancer (NCT03033225)^28,29^. Promising results from Phase I/II trials have shown significant PDT-induced necrosis, post-therapy^29^. One challenge with Visudyne® PDT, however, is non-selective localization of PSs in both cancerous and healthy tissue that can lead to off-target effects. For this reason, molecular targeted PDT offers tumor tissue specificity, thus overcoming challenges associated with non-specific and dose-limiting toxicity in cancer therapy^6,23,24,30-33^. Photoimmunoconjugates (PICs), are PSs covalently linked to antibodies^24,34^ and are known to confer specificity to otherwise non-specific photosensitizers. PIT is a molecular targeted PDT-based modality whereby photoactivation of PICs leads to receptor-specific tumor tissue phototoxicity^24^. PIT was pioneered by Levy and colleagues in 1983^35^ and was further developed by our team and several others^14,30,32-34,36,37^. PIT is emerging in the clinic, with a recent approval in Japan for Head and Neck Cancer and active Phase III trials in the US (NCT03769506) for patients with recurrent Head and Neck Squamous Cell Carcinoma. Considering the importance of Epidermal growth factor receptor (EGFR) in the targeted treatment and imaging of pancreatic and other cancers, it will be the target for PIT in our study^38-40^.

In this study, heterotypic 3D *in vitro* PDAC tumor models were developed with varying tumor cells and fibroblast ratios to recapitulate the cellular heterogeneity observed in the PDAC TME. These models were then used to explore the efficacy of clinically emergent EGFR-targeted PIT with respect to the formulation Visudyne® that is currently in clinical trials for PDAC patients. Furthermore, the mechanistic basis for the differences in therapeutic efficacy are explored and discussed in the context of stroma-induced resistance.

## Materials and Methods

### PIC synthesis

Benzoporphyrin derivative monoacid A (BPD, verteporfin) was conjugated to Cetuximab (Cet; Erbitux), obtained from Eli Lilly and Co. (Indianapolis, IN), according to a protocol established previously^30,32,33^. Briefly, Cetuximab (4 mg) was reacted overnight with Methoxy PEG Succinimidyl Carbonate, (MW 10000) (A3083, JenKem Technology, Plano, TX), dissolved in dimethyl sulfoxide (DMSO) at a 1:1 molar ratio. N-hydroxysuccinimidyl ester of BPD, prepared previously, was then reacted with the pegylated Cetuximab in a molar ratio of 9:1^33^. The reaction was carried out for 4 h at room temperature and the DMSO content was maintained at 40%. The crude product was centrifuged at 15 000 x *g* for 10 min to remove any aggregates, followed by purification using a Zeba spin desalting column (7K MWCO, 10 mL) (89893, Thermo Fisher Scientific, Waltham, MA). The eluent containing the purified PICs was then subjected to buffer exchange (5% DMSO in PBS) and concentrated using a 30 kDa NMWL Amicon® Ultra-15 Centrifugal Filter Unit (UFC903024, MilliporeSigma, Burlington, MA). The PICs were finally stored at 4 °C (in 5% DMSO) and remained stable for several months.

### Characterization of photosensitizer (PS) formulations - PIC and Visudyne®

PICs and Visudyne® (liposomal BPD, Bausch + Lomb) were diluted in DMSO and the BPD concentration was estimated using an Evolution™ 300 UV-Vis Spectrophotometer (ThermoFisher Scientific, Waltham, MA) and the known molar extinction coefficient of BPD at 687 nm; Ɛ_687_ = 34 895 M^-1^ cm^-1^. The concentration of Cetuximab was calculated using a Pierce™ BCA Protein Assay Kit (Thermo Fisher Scientific, Waltham, MA).

### Cell culturing and generation of fluorescent cell lines

Human Pancreatic Ductal Adenocarcinoma Cell line Mia PaCa-2 was obtained from American Type Culture Collection (ATCC) (CRM-CRL-1420™, Manassas, VA) and the patient-derived PCAF were a kind gift from Dr. Diane Simeone^41^. All cells were cultured in DMEM media supplemented with an antibiotic mixture containing Penicillin (100 I.U mL^-1^) and streptomycin (100 μg mL^-1^) (30-001-Cl, Corning, Corning, NY), and 10% heat-inactivated fetal bovine serum (FBS) (SH30071.03HI, Hyclone™, Marlborough, MA). The mCherry and EGFP expressing variants of Mia PaCa-2 cells and PCAFs, respectively, were generated by a previously established protocol^31,42^. Briefly, third-generation mCherry transfer plasmid pLV-mCherry (36084, Addgene, Watertown, MA) or the third-generation EGFP transfer plasmid pHAGE-CMV-EGFP-W (EvNO00061634, Harvard Plasmid Repository) were mixed with viral envelope plasmid pMD2.G (12259, Addgene, Watertown, MA) and viral packaging plasmid psPAX2 (12260, Addgene, Watertown, MA) at a molar ratio of 2:1:1 and transfected into the Lenti-X™ 293T packaging cell line (632180, TaKaRa Bio Inc., Kusatsu, Shiga, Japan) using Xfect™ Transfection Reagent (631317, TaKaRa Bio Inc., Kusatsu, Shiga, Japan) following manufacturer’s instruction. The viral supernatant was collected 48 h post-transfection, filtered using a 0.45 µm Puradisc 25 mm polyethersulfone Syringe Filter (6780-2504, GE Healthcare Life Sciences, Whatman™, Chicago, IL) to remove cellular debris, and concentrated using a Lenti-X™ Concentrator (631231, TaKaRa Bio Inc., Kusatsu, Shiga, Japan). Mia PaCa-2 cells and PCAFs were then transduced with the respective viral suspension at different multiplicity of infection (MOI) in the presence of 6 µg mL^-1^ hexadimethrine bromide (polybrene, Sigma H9268). Cells were sorted for mCherry and EGFP fluorescence using a BD FACSAria™ Cell Sorter (BD Biosciences, San Jose, CA), after 48 h of transduction. The sorted cells were expanded and cryopreserved in fetal bovine serum (10437028, Gibco™, ThermoFisher Scientific, Waltham, MA) with 8% DMSO. All cell lines used in this study tested negative for Mycoplasma contamination via MycoAlert™ PLUS Mycoplasma Detection Kit (LT07-710, Lonza, Basel, Switzerland).

### PDT dose-dependent toxicity in 3D MIA PaCa-2 spheroids

MIA PaCa-2 cells, cultured in 75 mm flasks, were trypsinized, counted and seeded in Corning Costar Ultra-Low attachment 96 well plates. The 3D spheroids were treated with PIC or Visudyne® for 24 h and 90 min, respectively, at a final PS concentration of 1 µM. In both cases, the photosensitizer containing culture media was removed, the spheroids washed thrice with PS-free culture media and irradiated, following previously described methods^43^. Briefly, 690 nm light was administered vertically through the plate bottom via a diode laser source at 150 mW cm^-2^ irradiance. The total energy deposit (light dose) was given at 0, 1, 2.5, 5, 7.5, 10, 15, 20, 25, 30 and 40 J cm^-2^. For 3D cultures, the culture plates were incubated after light administration for 3 days to allow complete development of cell death processes. The plates were imaged daily post-PDT with an Operetta CLS high-content analysis system (Perkin Elmer, LIVE configuration) with the incubation chamber set to 37 °C and 5% CO_2_. EGFP (ex/em: 475/500-570 nm) and mCherry (ex/em: 550/570-650 nm) fluorescence was collected via LED excitation (at 475 and 550 nm, respectively) and either a 5X, 0.16 NA air objective or a 10X, 0.3 NA air objective. 3D volumes were imaged in a z-stack format. Two fluorescence plus one brightfield z-stack were collected for each well using Harmony 4.6 (Perkin Elmer), an automated plate reading software. The full z-stack was reduced to a 2D image via maximum intensity projection (MIP) to analyze in-focus light. Images were analyzed via custom Image J script that applies the following analysis scheme to each well from each fluorescence channel to generate fluorescence volumetric data. The images were converted and converted to an 8-bit image and subjected to a background subtraction using a rolling ball algorithm with pixel size of 25. Equal brightness/contrast enhancement was performed to increase the histogram binning. Finally, the mean fluorescence of the corrected histograms was computed and used as a measure of cell viability.

### Simulating desmoplasia in vitro

MIA PaCa-2 cells and PCAFs were cultured as described earlier. For generating 3D spheroids, the Mia PaCa-2 cells and PCAFs were seeded in CellCarrier Spheroid ULA 96-well Microplates (6055330, PerkinElmer, Waltham, MA) with 0%, 10%, 25%, 50%, 75% and 90% PCAFs. The total number of cells were kept constant at 5000 cells per well. Spheroid size analysis and relative cellular ratios were calculated using Image J software. Quantification of growth and fractional relative viability were calculated as described in the previous section.

### Evaluation of PDT dose-dependent toxicity in in vitro desmoplastic PDAC tumors

3D MIA PaCa-2 tumor spheroids were cultured in DMEM with 10% FBS in Corning Costar Ultra-Low attachment 96 well plates with 0%, 10%, 25%, 50%, 75% and 90% of patient-derived PCAFs. The spheroids were treated with PIC or Visudyne® for 24 h and 90 min, respectively at a final PS concentration of 1 µM. As described previously, 690 nm light was administered vertically through the plate bottom via a diode laser source at 150 mW cm^-2^ irradiance as described previously. The total energy deposit (light dose) was given at 5 and 20 J cm^-2^ for the Visudyne® and PIC treated spheroids, respectively. Cell viabilities were calculated from the fluorescence intensity of the spheroids, calculated as described earlier. Relative viabilities were calculated by dividing fluorescence intensity of Mia PaCa-2-mCherry and PCAF-EGFP from the treated spheroids by the intensity of the respective Mia PaCa-2-mCherry and PCAF-EGFP from the untreated control spheroids, on the same day. Fractional relative viabilities were calculated using the following equations

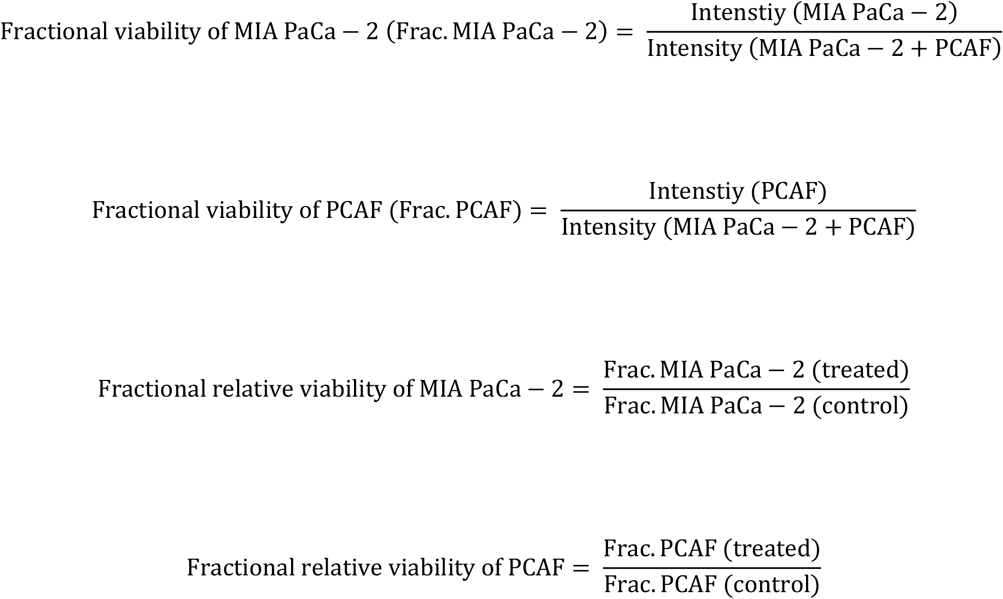

### Flow cytometry-based quantification of PS distribution in in vitro desmoplastic tumors

Heterotypic cultures (MIA PaCa-2 and PCAF with 100%, 50% and 25% MIA PaCa-2 cells) were grown in CellCarrier Spheroid ULA 96-well Microplates (6055330, PerkinElmer, Waltham, MA), as described previously. The spheroids were treated with PIC or Visudyne® for 24 h and 90 min, respectively at a final PS concentration of 1 µM. After incubation, spheroids from 12 wells with similar treatment condition were collected, pooled and centrifuged (5 min 500 *x g*). The spheroids were washed with once with 1 ml of 1x DPBS (21-031-CV, Corning®, Corning, NY) and resuspended in 0.25 ml of 0.05% Trypsin (25-052-CI, Corning®, Corning, NY). The spheroids were then incubated at 37 °C for a maximum of 3 min with mild agitation every 1 min. The detached cell suspensions were collected in DPBS containing 10% FBS and centrifuged for 5 min at 500 *x g*, the supernatant was aspirated and discarded, and the cells were redispersed in DPBS with 10% FBS. The cells were then transferred to flow cytometry tubes and temporarily stored on ice in the dark. For each condition, 10,000 cells were analyzed using a BD FACSAria II flow cytometer (BD Biosciences, San Jose, CA). Fluorescence of cell-associated BPD was measured using a 405 nm laser, a 655 nm long pass mirror and a 695/40 filter. Fluorescence from EGFP and mCherry were measured using 488 nm and 561 nm lasers, with 502 nm long pass mirror and 530/30 nm filter for EGFP and 600 nm long pass mirror and 610/20 nm filter.

### Photosensitizer distribution analysis in 3D spheroids (Confocal microscopy)

Mia PaCa-2 spheroids with and without PCAFs were generated as described earlier. The spheroids were then treated with PIC and Visudyne® at 1 µM BPD equivalent concentration for 90 min and 24 h, respectively. Following incubation, the spheroids were washed thrice with DPBS and fixed with 4% formalin for 15 min. Excess formalin was quenched with 1 M Glycine and the spheroids were stained with 1 µg/ml Hoechst 33342 solution (62249, ThermoFisher Scientific, Waltham, MA). The organoids were imaged using a 4x objective and appropriate lasers and filters for Hoechst, EGFP, mCherry and BPD. Across all organoids, Z-stack images were acquired to cover the entire depth of the spheroid. 2D cross-sections from the central plane (corresponding to approximately 50% depth) of each spheroid was selected to generate the profile plots and study BPD distribution and penetration.

## Results

### Photosensitizer formulations and drug-light interval

Visudyne® is a lipid-based formulation of Verteporfin with a hydrodynamic diameter of ~ 524.8 nm (Polydispersity index of 0.88)^27^. It has been previously used in several pre-clinical and clinical studies with a photosensitizer-light interval of 60 – 90 min^19,28,29^. It mostly localizes in the mitochondria and the ER^27^. The EGFR-targeted photosensitizer formulation: PIC was synthesized through a previously established protocol^32,33^. This protocol consistently results in the generation of PICs with 5-7 photosensitizer molecules per molecule of Cetuximab. The conjugate also has a PEG molecule which assists in solubilization of the antibody, preventing non-specific uptake and enhancing the *in vivo* half-life of the antibody^44,45^. The fluorescence from BPD within the PICs is usually quenched and can be recovered by PIC activation after the antibody is internalized in the target cells through receptor mediated endocytosis followed by lysosomal degradation of the conjugate^30,31^. Post degradation the hydrophobic PS can migrate to the Mitochondria and the ER resulting in the induction of apoptosis through the lysosomal and mitochondrial route^24,46,47^. PICs have been used previously in our lab *in vitro* and *in vivo* with a photosensitizer-light interval of 24 h^30^. Moreover, similar clinically approved antibody photosensitizer conjugates have been used with a photosensitizer-light interval of 24 h for PIT of head and neck cancers^48,49^. Therefore, in this study, the DLI for Visudyne® and PIC was fixed at 90 min and 24 h, respectively (**Table 1**).

**Table 1:**
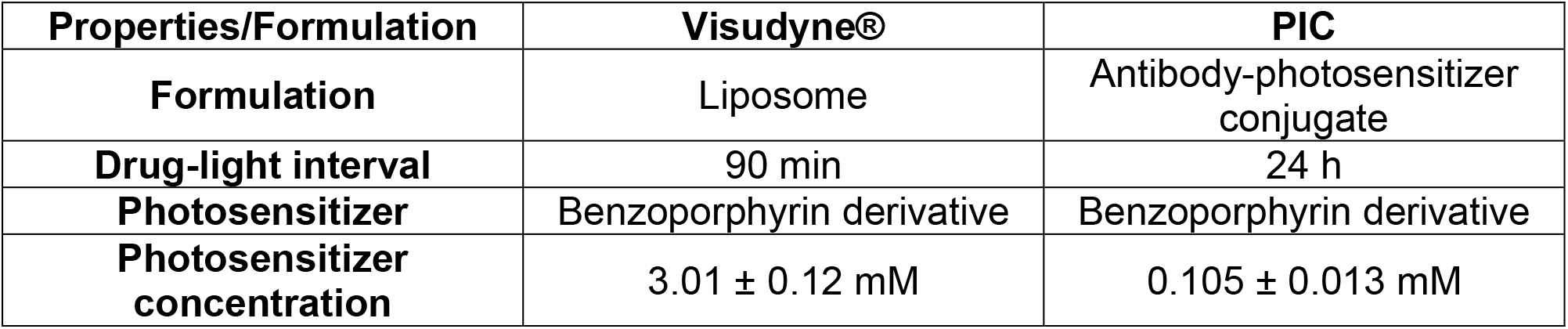
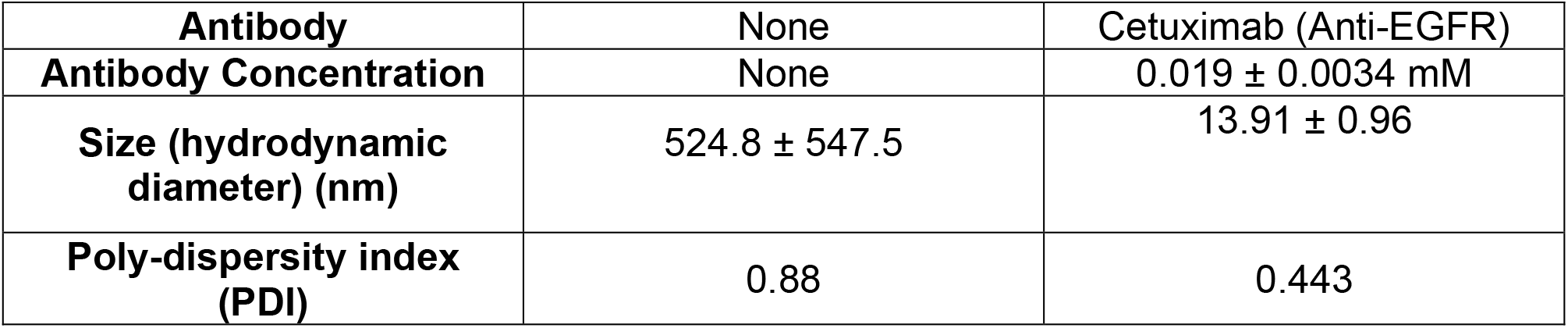
Physico-chemical characterization of Visudyne® and Photo-immunoconjugates (PICs):

| **Properties/Formulation** | **Visudyne®** | **PIC** |
| --- | --- | --- |
| **Formulation** | Liposome | Antibody-photosensitizer conjugate |
| **Drug-light interval** | 90 min | 24 h |
| **Photosensitizer** | Benzoporphyrin derivative | Benzoporphyrin derivative |
| **Photosensitizer concentration** | 3.01 ± 0.12 mM | 0.105 ± 0.013 mM |
| **Antibody** | None | Cetuximab (Anti-EGFR) |
| **Antibody Concentration** | None | 0.019 ± 0.0034 mM |
| **Size (hydrodynamic diameter) (nm)** | 524.8 ± 547.5 | 13.91 ± 0.96 |
| **Poly-dispersity index (PDI)** | 0.88 | 0.443 |

### Generation of MIA PaCa-2-mCherry and PCAF-EGFP cell lines

Tagging cell lines with fluorescent proteins is an elegant method to decipher the specificity of targeted/non-targeted agents in a heterocellular model^14,17^. For this reason, in this study we generated fluorescently tagged MIA PaCa-2 and PCAF lines to decipher the specificity of the PS formulations in a MIA PaCa-2/PCAF heterotypic 3D organoid. The Mia PaCa-2 and PCAF cell lines were transduced to express mCherry and EGFP, respectively. The fluorescence expression of the two cell lines was confirmed by flow cytometry and fluorescence imaging. As shown in **Figure 1**, Mia PaCa-2 and PCAF cells showed fluorescence for mCherry and EGFP, respectively.

**Figure 1:**
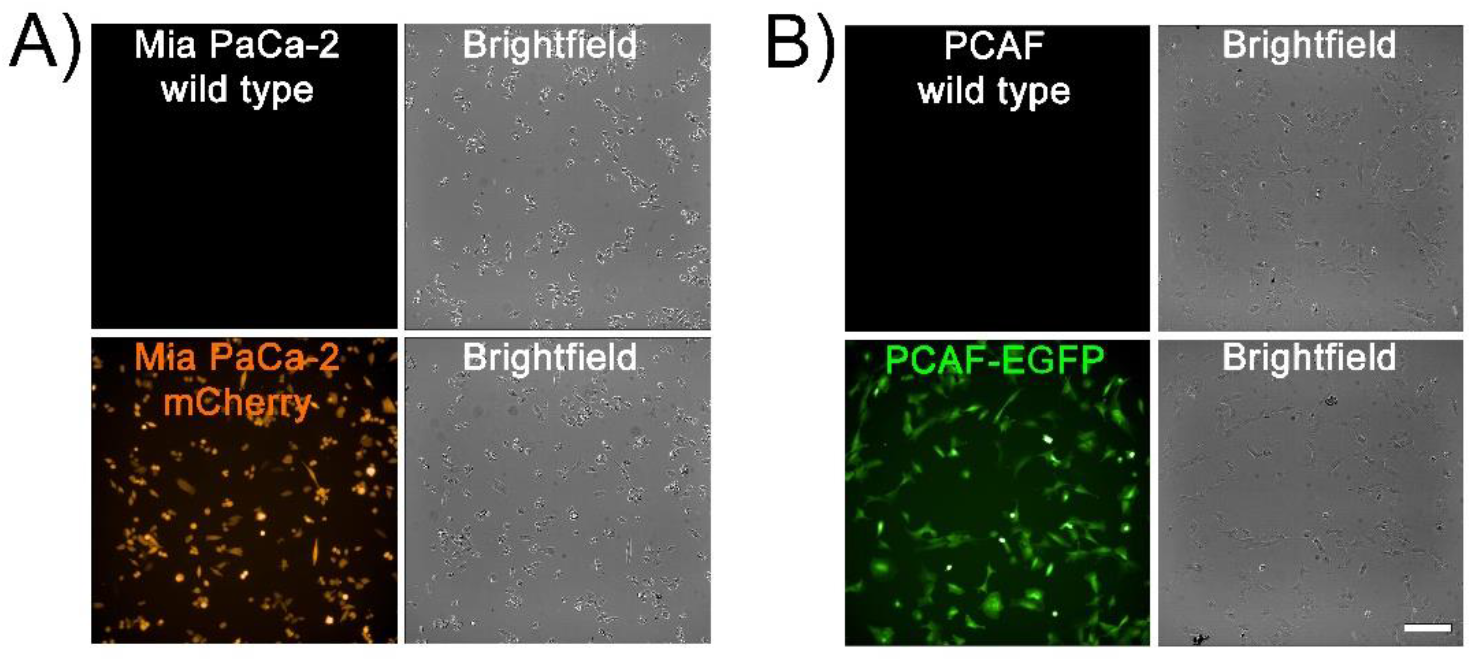
Representative images of 2D cultures of wild type (**A)** MIA PaCa-2 cells and Mia PaCa-2 cells expressing mCherry, (**B**) wild type PCAFs and PCAFs expressing EGFP. (scale bar = 200 µm).

### Generation of 3D spheroids and simulating desmoplasia

To generate different levels of stromal content often observed in PDAC and other tumors, Mia PaCa-2-mCherry were mixed with different PCAF percentage (0%, 10%, 25%, 50%, 75% and 90%), with the total cell numbers being fixed at 5000 cells. **Figure 2A** shows the morphological and fluorescent characteristics (MIA PaCa-2-mCherry and PCAF-EGFP, pseudo-colored as orange and green, respectively) of the spheroids formed with different PCAF percentage on day 2 after cell seeding. PCAFs were observed to grow in clusters in spheroids formed with low PCAF percentage (10 – 50%), while the ones with higher PCAF percentage (75 and 90%) showed a single cluster of PCAFs with interspersed MIA PaCa-2 cells. Longitudinal monitoring of the spheroids was performed over 5 days using an Operetta CLS High-Content Analysis System. **Figure 2B** shows the changes in the relative cellular proportion of the two cell lines monitored over time (5 days). The growth curves were generated by quantifying intensities of the two fluorophores, normalized over their respective intensities obtained at day 1. As shown, MIA PaCa-2 cells rapidly outnumbered the PCAFs over the 5-day period of culture, with PCAF percentage in the different spheroids on day 5 being 0.2%, 0.6%, 1.1%, 4% and 28.4% for the spheroid cultures initiated with 10%, 25%, 50%, 75% and 90% PCAFs, respectively. Previous studies have also suggested a decrease in fibroblast population when co-cultured with cancer cells or in cancer cell conditioned media^50^. **Figure 2C** shows the size variation of the spheroids with increasing PCAF percentage on day 2 after cell seeding. A decrease in spheroid diameter with increasing PCAF percentage was observed, with spheroids formed with 90% PCAF being the smallest (308.68 ± 21.15 µm) as compared to the ones with no PCAFs (1003.70 ± 73.55 µm). Spheroids formed with 50% MIA PaCa-2 and PCAFs were the most polydisperse characterized by a high standard deviation (714.84 ± 91.63 µm). A comparison of relative cellular ratios and spheroid sizes (on day 2 post-cell seeding) are depicted as pie charts (**Figure 2D**).

**Figure 2:**
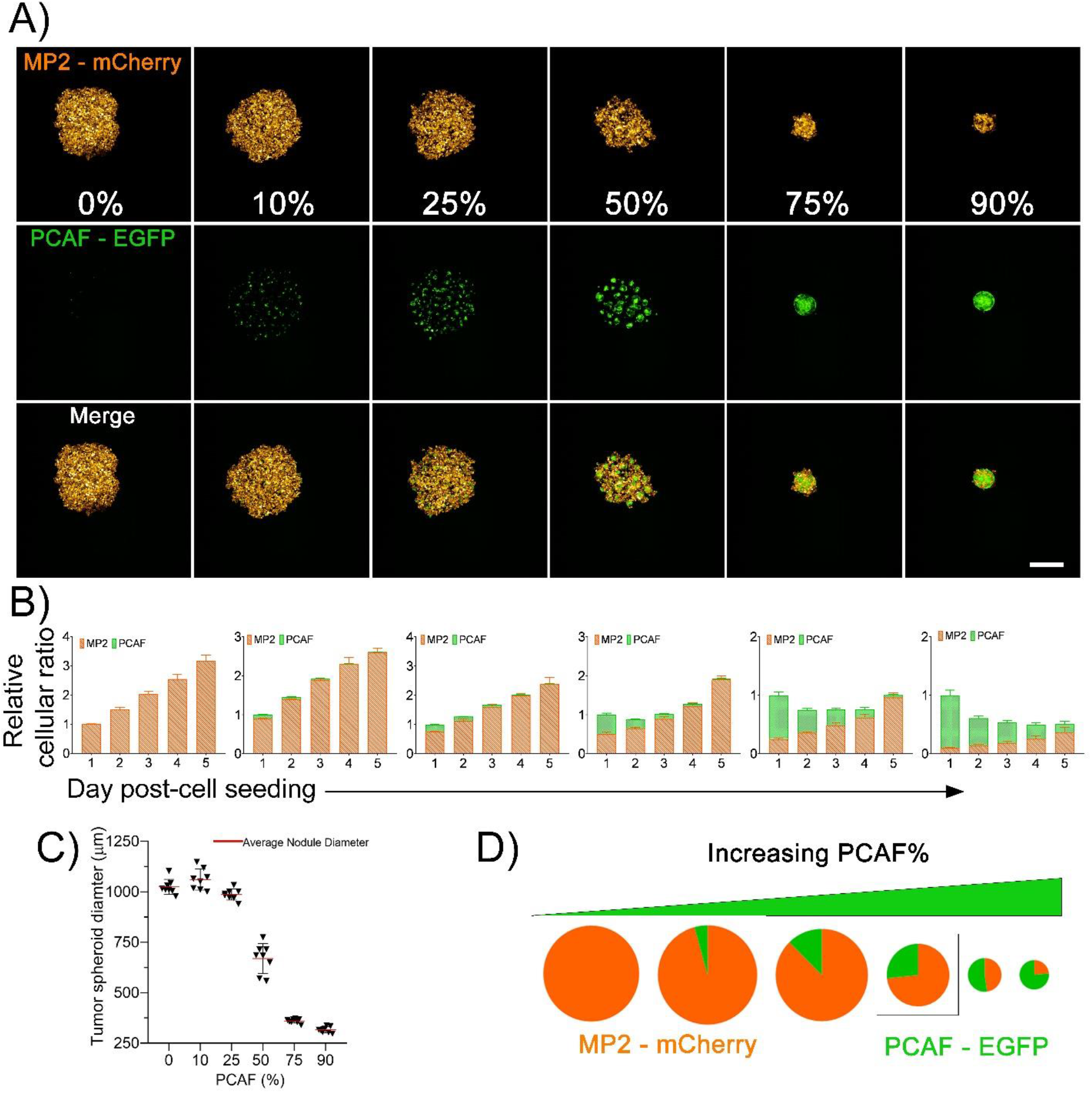
Characterization of 3D *in vitro* heterotypic cultures generated with different ratios of Mia PaCa-2 and PCAFs. (**A**) Maximum intensity projection (MIP) of fluorescence images of spheroids developed with increasing PCAF percentage. (left to right). Mia PaCa-2 are pseudo-colored as orange and PCAFs are pseudo-colored as green. Scale bar corresponds to 500 µm. (**B**) Relative cellular ratio of Mia PaCa-2 and PCAFs in the different spheroids (left to right, 0%, 10%, 25%, 50%, 75% and 90% PCAFs) as cultured for 5 days. The Mia PaCa-2 cells rapidly out-numbered the PCAFs during culture. The cellular ratios were calculated based on the 2D fluorescent MIP images for these spheroids using image J. Mia PaCa-2 are represented as orange bars and PCAFs are represented as green bars. (**C**) Diameter of the tumor spheroids formed with different PCAF percentage on day 2 of culture. (**D**) Pie charts providing a comparison of relative cellular ratio and spheroid size on day 2 post-cell seeding. Size of the pie charts represent the size of spheroids, while the orange and green colors represent the fraction of Mia PaCa-2 and PCAFs, respectively.

### Light dose dependent phototoxicity of Visudyne® and PIT on homotypic Mia PaCa-2 3D spheroids

Next, to compare the efficacy of Visudyne®-PDT and PIT in inducing cytotoxicity in 3D MIA PaCa-2 cultures, the two PS formulations were incubated for time periods corresponding to their clinical drug-light interval (90 min and 24 h, respectively), followed by irradiation at different light doses. **Figure 3A**, shows the experimental schedule for treatment of the spheroids and **Figure 3B** shows the mCherry fluorescence intensity of the MIA PaCa-2 spheroids, on day 3 post-PDT. As suggested by the fluorescence images, the mCherry intensity of the spheroids decreased with increasing light doses, suggesting a decrease in viability of the tumor spheroids. **Figure 3C** provides a quantitative analysis of the images shown in figure 3B. Visudyne®-PDT was found to be more effective, as compared to PIT, in inducing cytotoxicity even at lower light doses with a relative viability of 0.18 ± 0.07, 0.06 ± 0.01 and 0.04 ± 0.01 at 1, 2.5 and 5 J/cm^2^, respectively. As compared to Visudyne®-PDT, PIT resulted in a gradual decrease in relative viability with increasing light dose and showed a minimal decrease in relative viability at 1, 2.5 and 5 J/cm^2^ (1.00 ± 0.08, 0.94 ± 0.40 and 0.79 ± 0.24, respectively). A further increase in light dose to 7.5, 10, 15 and 20 J/cm^2^, resulted in a relative viability of 0.42 ± 0.21, 0.28 ± 0.16, 0.10 ± 0.04 and 0.066 ± 0.03, respectively. As the relative viability of PIT at 20 J/cm^2^ was comparable to the relative viability of Visudyne®-PDT at 5 J/cm^2^, these doses were used subsequently to evaluate the efficacy of PDT using the two formulations in spheroids formed with different PCAF percentage for simulating stromal content heterogeneity.

**Figure 3:**
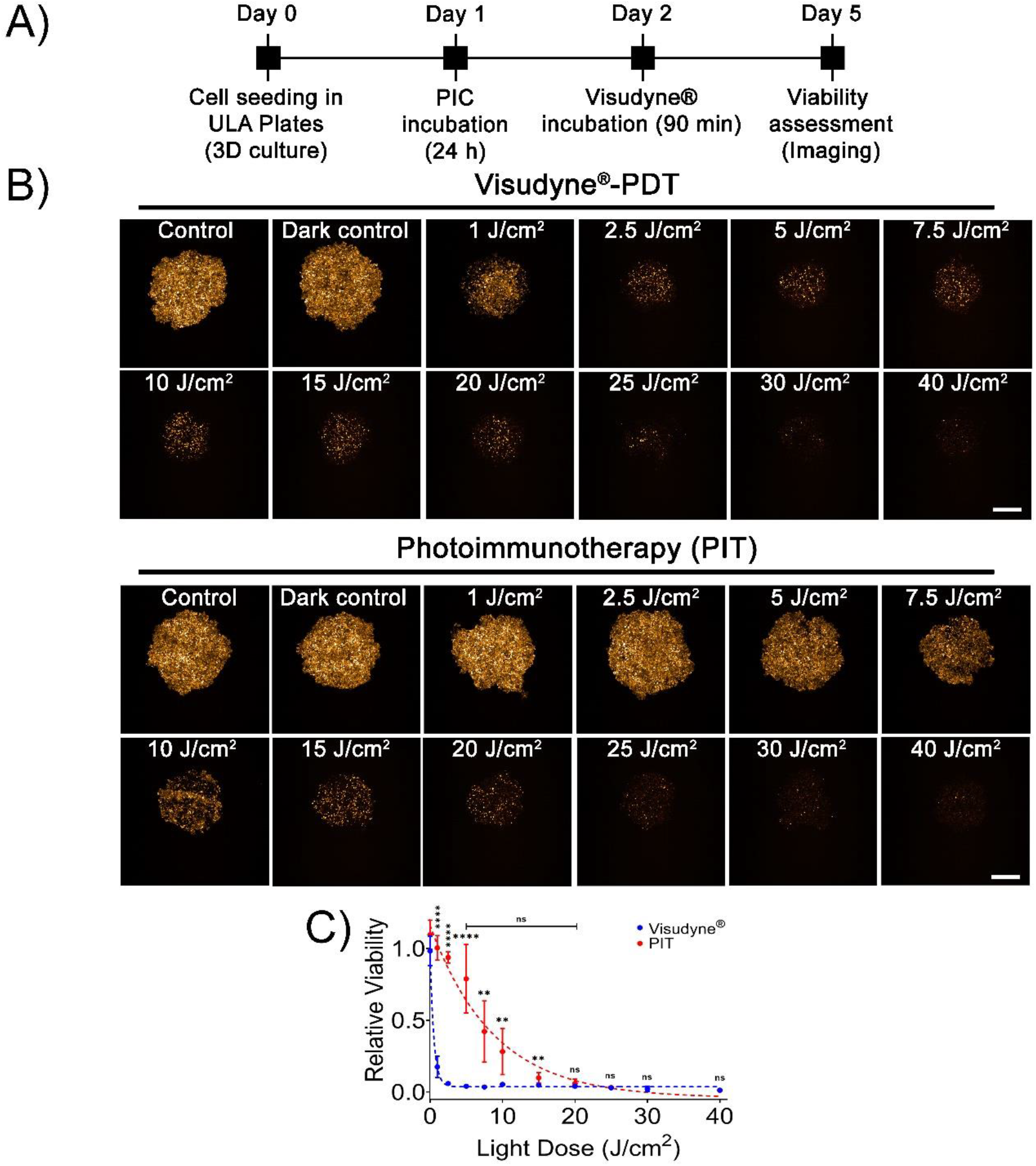
Dose-dependent response of Mia PaCa-2 homotypic 3D cultures to Visudyne®-PDT and PIT. (**A**) Experimental schedule for Visudyne® and PIC-PDT. Cells were seeded in CellCarrier Spheroid ULA 96-well Microplates followed by incubation with either PICs or Visudyne® for 24 h and 90 min, respectively. The spheroids were then irradiated with different light doses and imaged for 3 days post-PDT. (**B**) Maximum intensity projection (MIP) of fluorescence images of MP2 spheroids on day 3 post-PDT. Scale bar corresponds to 500 µm. (**C**) Quantitative analysis of mCherry fluorescence from spheroids treated with PDT. PIT resulted in a gradual decrease in viability, while Visudyne®-PDT resulted in a sudden drop in viability with increasing light dose. Data points were fit in a non-linear exponential decay equation (R^2^ = 0.98 and 0.91 for Visudyne®-PDT and PIT, respectively). (Data points for Visudyne®-PDT and PIT are represented in blue and red dots, while the non-linear fits are represented as dotted lines of the same color). Data are presented as mean ± S.D (n ≥8), analyzed using Welch’s t-test analysis. P-values < 0.05 were considered to be significant and are indicated by asterisks as follows: nsP>0.05, *P<0.05, **P<0.01, ***P<0.001 and ****P<0.0001.

### Efficacy of and Visudyne®-PDT and PIT on heterotypic 3D spheroids: Impact of stromal content

Next, the efficacy of Visudyne®-PDT and PIT was evaluated on Mia PaCa-2-PCAF heterocellular cultures, generated with varying Mia PaCa-2 to PCAF ratios, using the light dose (5 J/cm^2^ and 20 J/cm^2^ for Visudyne®-PDT and PIT, respectively) and drug light intervals (90 min and 24 h for Visudyne®-PDT and PIT, respectively) standardized for monotypic MIA PaCa-2 cultures. **Figure 4** shows representative images of heterotypic spheroids treated with Visudyne®-PDT and PIT. Control spheroids are shown in the top panel for comparison. As evident from the images, PIT appeared to be more efficient than Visudyne®-PDT in inducing phototoxicity as the PCAF percentage increased. In contrary, Visudyne®-PDT was effective only in spheroids with low PCAF percentage and its efficacy decreased as the percentage of PCAF increased in the spheroids.

**Figure 4:**
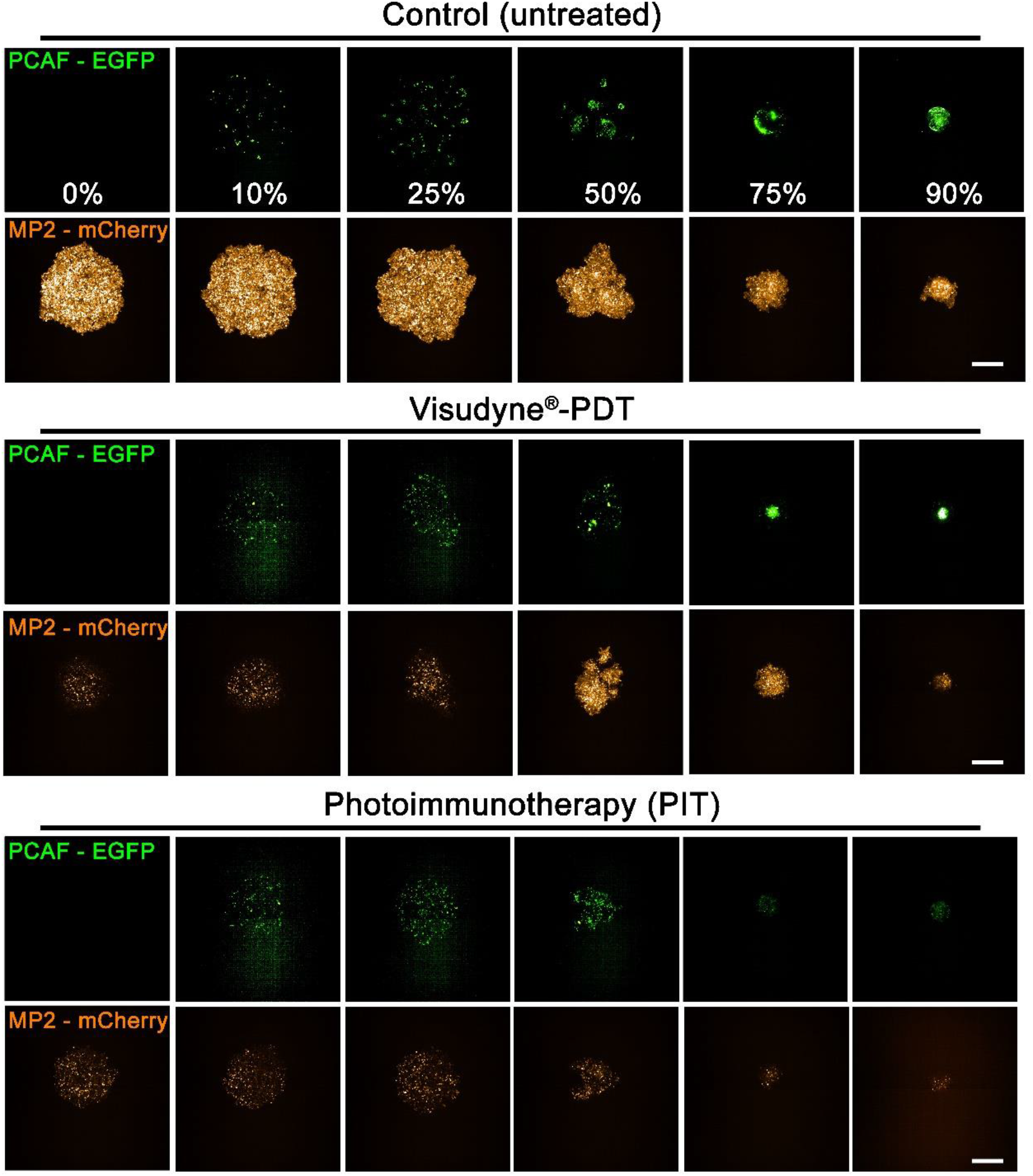
Response of heterotypic Mia PaCa-2-PCAF 3D cultures to Visudyne®-PDT and PIT. Maximum intensity projection (MIP) of fluorescence signals from Mia PaCa-2-PCAF spheroids with different PCAF percentage on day 3 post-PDT. PCAF percentage is mentioned in the top panel (control spheroids PCAF-EGFP panel). Scale bar corresponds to 500 µm.

Figure 5. shows a quantitative analysis of images in **Figure 4**. Relative viabilities were calculated by dividing fluorescence intensity of Mia PaCa-2-mCherry and PCAF-EGFP from the treated spheroids by the intensity of the respective Mia PaCa-2-mCherry and PCAF-EGFP from the untreated control spheroids, on the same day. As evident, from **figure 5B and 5C**, at low PCAF percentage (0%-25%) both Visudyne®-PDT and PIT were equally effective in significantly reducing the relative viability of Mia PaCa-2 cells and PCAFs in these heterotypic cultures (**red graphs in Figure 5B for Mia PaCa-2 and 5C for PCAF**). However, increasing the PCAF percentage to 50% and beyond (75% and 90%) resulted in a decrease in PDT efficacy of Visudyne®, with the relative viability of MIA PaCa-2 cells being 0.32 ± 0.21, 0.57 ± 0.20 and 0.42 ± 0.13. Interestingly, the relative viability of PCAFs under these conditions (50%, 75% and 90% PCAF treated with Visudyne®-PDT) increased significantly, even higher than the corresponding controls; 0.27 ± 0.21, 1.19 ± 0.60 and 1.78 ± 0.7 for spheroids formed with 50%, 75% and 90% PCAFs, respectively. PIT however was not affected by PCAF percentage and maintained its potent phototoxicity even in the spheroids with high PCAF percentage. The relative viability of spheroids treated with PIT was 0.032 ± 0.02, 0.03 ± 0.03, and 0.03 ± 0.01 for Mia PaCa-2 and 0.05 ± 0.03, 0.02 ± 0.01 and 0.01 ± 0.00 for PCAFs in spheroids with 50%, 75% and 90% PCAF percentage (**blue graphs in Figure 5B for Mia PaCa-2 and 5C for PCAF**). In comparison, PIT was 10 folds, 19 folds and 14 folds more phototoxic in spheroids with 50%, 75% and 90% PCAFs, respectively, as compared to Visudyne®-PDT. **Figure 5D** and **5E** show the relative fraction of the surviving population in the spheroids treated with PIT (**Figure 5D**) and those treated with Visudyne®-PDT (**Figure 5E**). As suggested by the orange bars in **Figure 5D**, PIT reduced the relative viability of Mia PaCa-2 significantly and to a similar extent, irrespective of the PCAF percentage. However, the PCAFs were less effected by PIT in the spheroids with 10%, 25% and 50% PCAFs, which corresponds to the lower EGFR expression by the PCAF cells with respect to the Mia PaCa-2 cells^23^. In contrast, Visudyne® effectively reduced the relative fraction of the surviving population for both Mia PaCa-2 and PCAFs in spheroids with 0%, 10% and 25% PCAFs. However, in spheroids with 50%, 75% and 90% PCAFs, Visudyne® was less effective in reducing the viability of Mia PaCa-2 cells and PCAFs in which case it promoted growth. This observation in of itself is highly intriguing, as the results suggest that sub-therapeutic stress to the heterotypic spheroids with high PCAF percentage specifically results in a stroma-dependent pro-survival response that may increase the aggressiveness of the disease.

**Figure 5:**
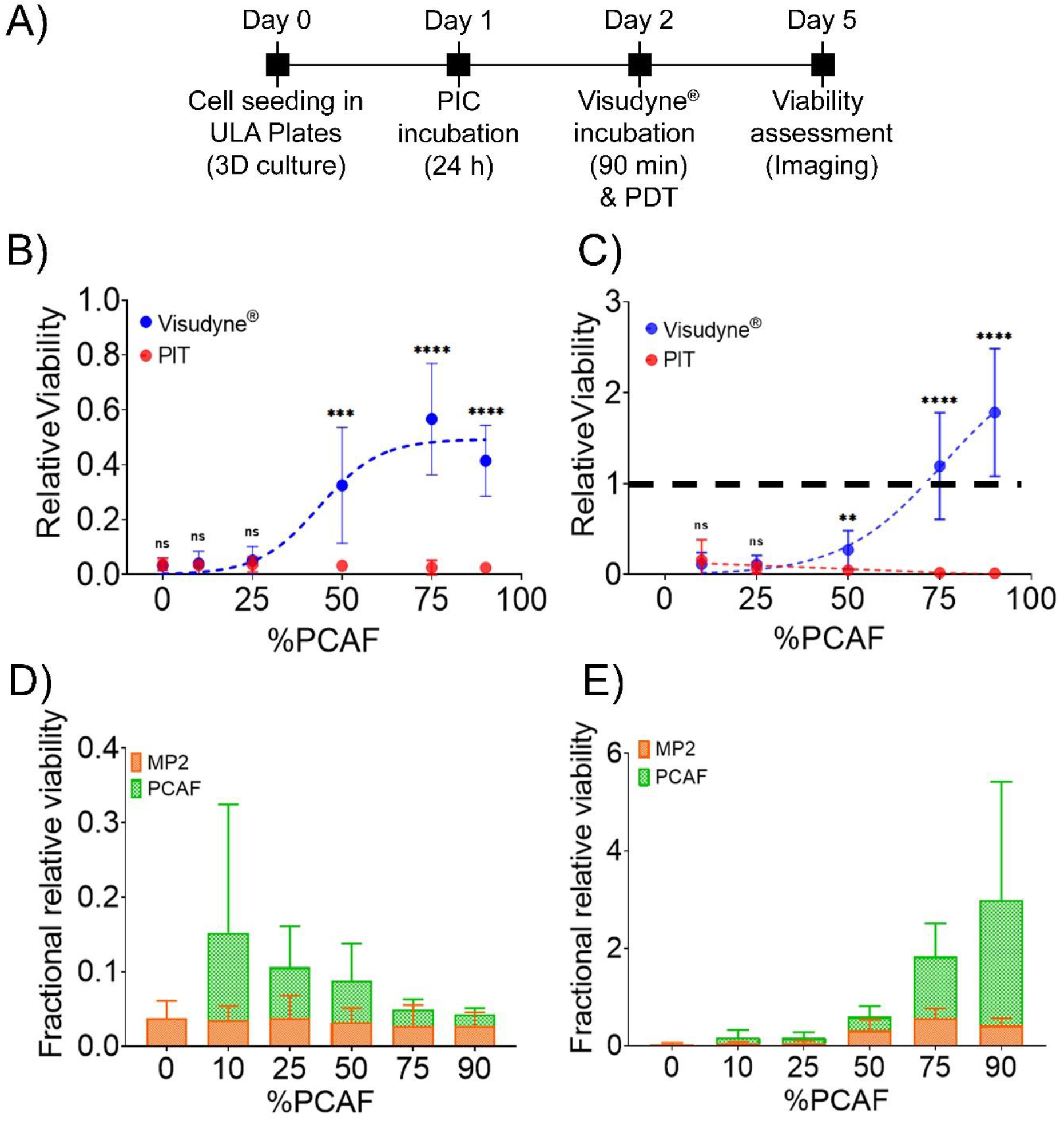
Quantitation of cell viability in heterotypic Mia PaCa-2-PCAF 3D cultures in response to Visudyne®-PDT and PIT. (**A**) Experimental schedule for Visudyne®-PDT and PIT of heterotypic Mia PaCa-2-PCAF spheroids. Cells were seeded in CellCarrier Spheroid ULA 96-well Microplates followed by incubation with either Visudyne® or PIC for 90 min and 24 h, respectively. The spheroids were then irradiated with a light dose of 5 J/cm^2^ (for Visudyne® treated spheroids) and 20 J/cm^2^ (for PIC treated spheroids) followed by imaging for 3 days. Relative viability of individual Mia PaCa-2 (**B**) and PCAF (**C**) populations in spheroids (formed with different PCAF percentage) treated with either PIT (red circles) or Visudyne®-PDT (blue circles). Both Visudyne®-PDT and PIT reduced the relatively viability of Mia PaCa-2 and PCAFs significantly in spheroids with low PCAF percentage (0%, 10% and 25%). However, increasing the PCAF percentage further resulted in a decrease in efficiency of Visudyne®-PDT, while PIT was relatively unaffected. Relative viabilities were calculated by dividing fluorescence intensity of Mia PaCa-2-mCherry and PCAF-EGFP from the treated spheroids by the intensity of the respective Mia PaCa-2-mCherry and PCAF-EGFP from the untreated control spheroids, on the same day. Data are presented as mean ± S.D (n ≥3), analyzed using Welch’s t-test analysis. P-values < 0.05 were considered to be significant and are indicated by asterisks as follows: ^ns^P>0.05, *P<0.05, **P<0.01, ***P<0.001 and ****P<0.0001. (**C**) Fractional relative viability of Mia PaCa-2 and PCAFs in spheroids treated with PIT (**D**) and Visudyne®-PDT (**E**) calculated as described in the methods. PIT was relatively more phototoxic (although not statistically significant) to Mia PaCa-2 cells in spheroids formed with low PCAF percentage (10%, 25% and 50%). However, increasing the PCAF percentage resulted in a decrease in selectivity and in spheroids with 75% and 90% PCAFs, the fractional relative viability of Mia Paca-2 and PCAFs were similar. On the other hand, spheroids treated with Visudyne®-PDT showed an increase in fractional relative viability of PCAFs in spheroids with 75% and 90% PCAFs. Mia PaCa-2 are represented as orange bars and PCAFs are represented as green bars. Data are presented as mean ± S.D (n ≥3).

### Evaluation of photosensitizer distribution in Mia PaCa-2 and PCAFs in spheroids with different levels of stroma

To gain mechanistic insights into the differences in efficacy of the two photosensitizer formulations with increasing desmoplasia, we studied the distribution profile of the photosensitizer delivered through Visudyne® and PIC and quantified the PS uptake. Flow cytometry analysis on tumor spheroids formed with 0%, 50% and 75% PCAFs, after treatment with the two photosensitizers was performed as described in the experimental scheme in **Figure 6A**. As shown in **Figure 6B**, irrespective of the PCAF percentage, the median BPD fluorescence for spheroids treated with PIC was significantly lower as compared to the ones treated with Visudyne®, suggesting a higher PS uptake in Visudyne® treated spheroids. Interestingly, the PS uptake in Mia PaCa-2 cells was similar in spheroids with different PCAF percentage (different desmoplasia) when delivered through PICs (Median BPD fluorescence 879 ± 87.2, 830.25 ± 132.57 and 778.25 ± 120.00 for spheroids with 0%, 50% and 75% PCAFs, respectively). The median BPD fluorescence for the corresponding PCAFs in these spheroids was slightly lower; 435.5 ± 100.37 and 379.75 ± 91.63 for spheroids with 50% and 75% PCAFs respectively). In contrast, the BPD uptake in Mia PaCa-2 cells when delivered through Visudyne® decreased with increasing PCAF percentage (increasing desmoplasia) (Median BPD fluorescence 6441 ± 597.34, 4368 ± 211.42 and 2578.5 ± 405.17 for spheroids with 0%, 50% and 75% PCAFs, respectively). The median BPD fluorescence for the corresponding PCAFs in these spheroids was lower; 3413.5 ± 58.70 and 2102.5 ± 231.22 for spheroids with 50% and 75% PCAFs, respectively). To study the distribution profile (heterogeneity/homogeneity) of BPD, we plotted contour plots of BPD fluorescence for the different spheroids treated the two formulations. As shown in **figure 6C**, Mia PaCa-2 only spheroids showed a homogeneous distribution of BPD fluorescence as indicated by narrow contour plots for both Visudyne® and PICs. Importantly, increasing PCAF percentage (desmoplasia) did not have a significant effect on BPD distribution when delivered through PICs, as evident by the narrow contour plots for BPD in Mia PaCa-2 cells in these spheroids (**Figure 6C, red contour plots in upper panel**). However, the BPD distribution profiles in Mia PaCa-2 cells, when delivered through Visudyne®, appeared to be broad, suggesting a heterogeneous distribution in the spheroids with high PCAF percentage (**Figure 6C, blue contour plots in upper panel**). The BPD distribution profiles in the corresponding PCAFs for the same spheroids incubated with Visudyne® also had broad contour plots (**Figure 6C, blue contour plots in lower panel**), while the ones treated with PICs had narrow contour plots, suggesting a comparatively homogenous distribution (**Figure 6C, red contour plots in lower panel**).

**Figure 6:**
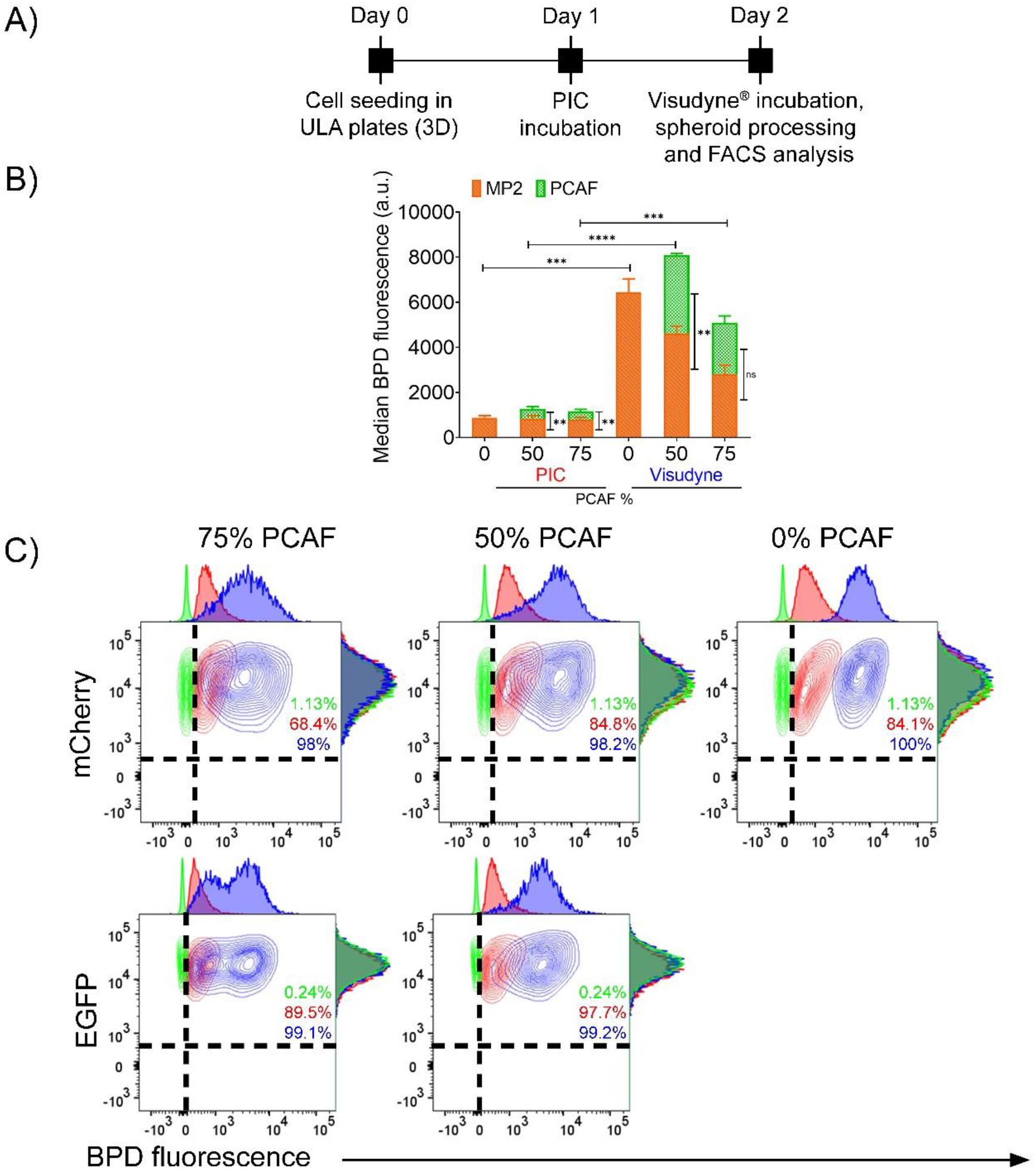
Photosensitizer distribution in heterotypic 3D spheroids. (**A**) Experimental timeline for studying BPD uptake in homotypic and heterotypic spheroids using Visudyne® and PIC. Cells were seeded in CellCarrier Spheroid ULA 96-well Microplates followed by incubation with either PICs or Visudyne® for 24 h and 90 min, respectively. The spheroids were then washed, trypsinized and analyzed through flow cytometry. (**B**) Median BPD fluorescence of spheroids treated with PICs and Visudyne®. Green bars represent PCAFs while orange bars represent MP2 cells. BPD distribution in Mia PaCa-2 and PCAFs were unaffected by PCAF percentage in the spheroids and remained same. However, BPD uptake in Mia-PaCa-2 cells when delivered through Visudyne® was significantly reduced with increasing PCAF percentage. The corresponding BPD uptake in PCAFs of these spheroids did not change significantly with increasing PCAF percentage. Mia PaCa-2 are represented as orange bars and PCAFs are represented as green bars. Data are presented as mean ± S.D (n ≥3), analyzed using Welch’s t-test analysis. P-values < 0.05 were considered to be significant and are indicated by asterisks as follows: ^ns^P>0.05, *P<0.05, **P<0.01, ***P<0.001 and ****P<0.0001. (**C**) Contour plots for analyzing BPD distribution heterogeneity in homotypic and heterotypic spheroids. Mia PaCa-2 cells (upper panel) and PCAFs (lower panel) in spheroids with 75% and 50% PCAFs showed broader contour plots for Visudyne® (blue), while the contour plots for PIC (red) were narrow, suggesting a homogeneous distribution of BPD after PIC treatment.

### Photosensitizer distribution profile in MP2 and PCAF spheroids with different levels of stroma

To gain further insights into the distribution profile of the photosensitizer (BPD) in the spheroids with different levels of desmoplasia, the spheroids were cultured and treated with either PIC or Visudyne® as shown in the experimental scheme (**Figure 7A and 8A**). Confocal fluorescence microscopy was performed, and optical sections were obtained at depths of 10 µm. **Figure 7B** shows the projection of the central section of the different spheroids (different PCAF percentage) treated with PICs. Profile plots were generated to analyze BPD fluorescence across the central section of these spheroids. Profile plots for the spheroids treated with PIC showed homogenous distribution irrespective of the PCAF percentage. On the contrary, while spheroids with 0%, 10%, and 25% PCAF, when treated with Visudyne® showed a homogeneous distribution of BPD as suggested by the profile plots. In contrast, spheroids with 50%, 75% and 90% PCAFs showed a higher accumulation of BPD in the peripheral layers and a lower accumulation in the center (**Figure 8**). These results correlate with the heterogeneous BPD distribution observed in the flow cytometry analysis where homogenous distribution (narrow contour plots) was observed in spheroids with 0% PCAF, while heterogeneous distribution (broader contour plots) was observed in spheroids with 50% and 75% PCAFs.

**Figure 7:**
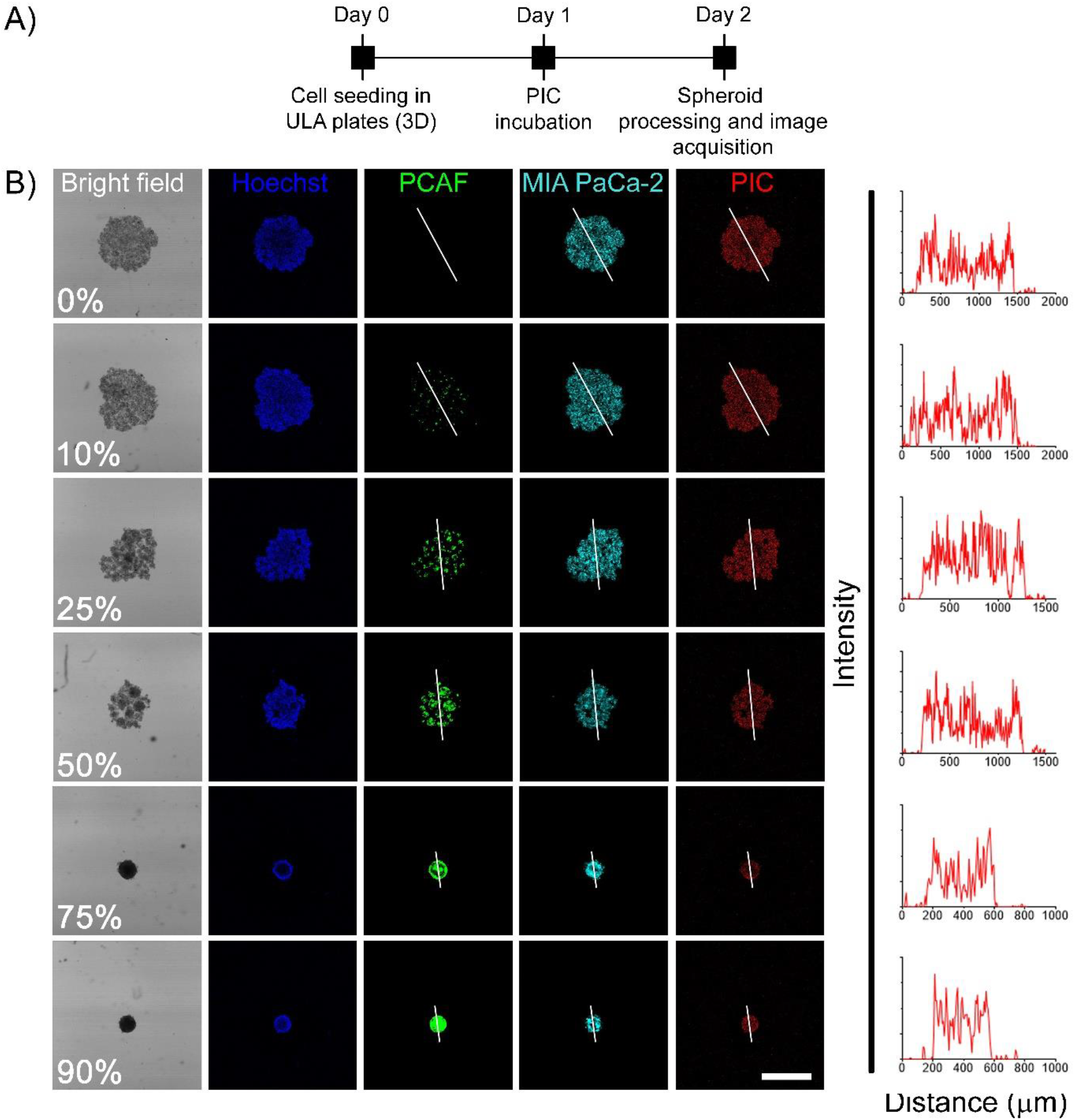
PIC administered photosensitizer distribution in heterotypic 3D spheroids. (**A**) Experimental timeline for studying BPD uptake and distribution in homotypic and heterotypic spheroids using PICs. Cells were seeded in CellCarrier Spheroid ULA 96-well Microplates followed by incubation with PICs for 24 h. The spheroids were then washed, fixed in PFA, counter-stained with Hoechst and visualized under a confocal fluorescence microscope. (**B**) Representative confocal images of the central optical section of the 3D spheroids. Mia PaCa-2 cells and PCAFs are pseudo-colored as cyan and green. Nuclei are stained with Hoechst (pseudo-colored as blue) and BPD is shown in red. Profile plots were generated using image J to monitor BPD distribution across the central plane. As suggested by the profile-plots, the distribution of BPD was homogeneous across the central section irrespective of the spheroid composition (scale bar = 500 µm).

**Figure 8:**
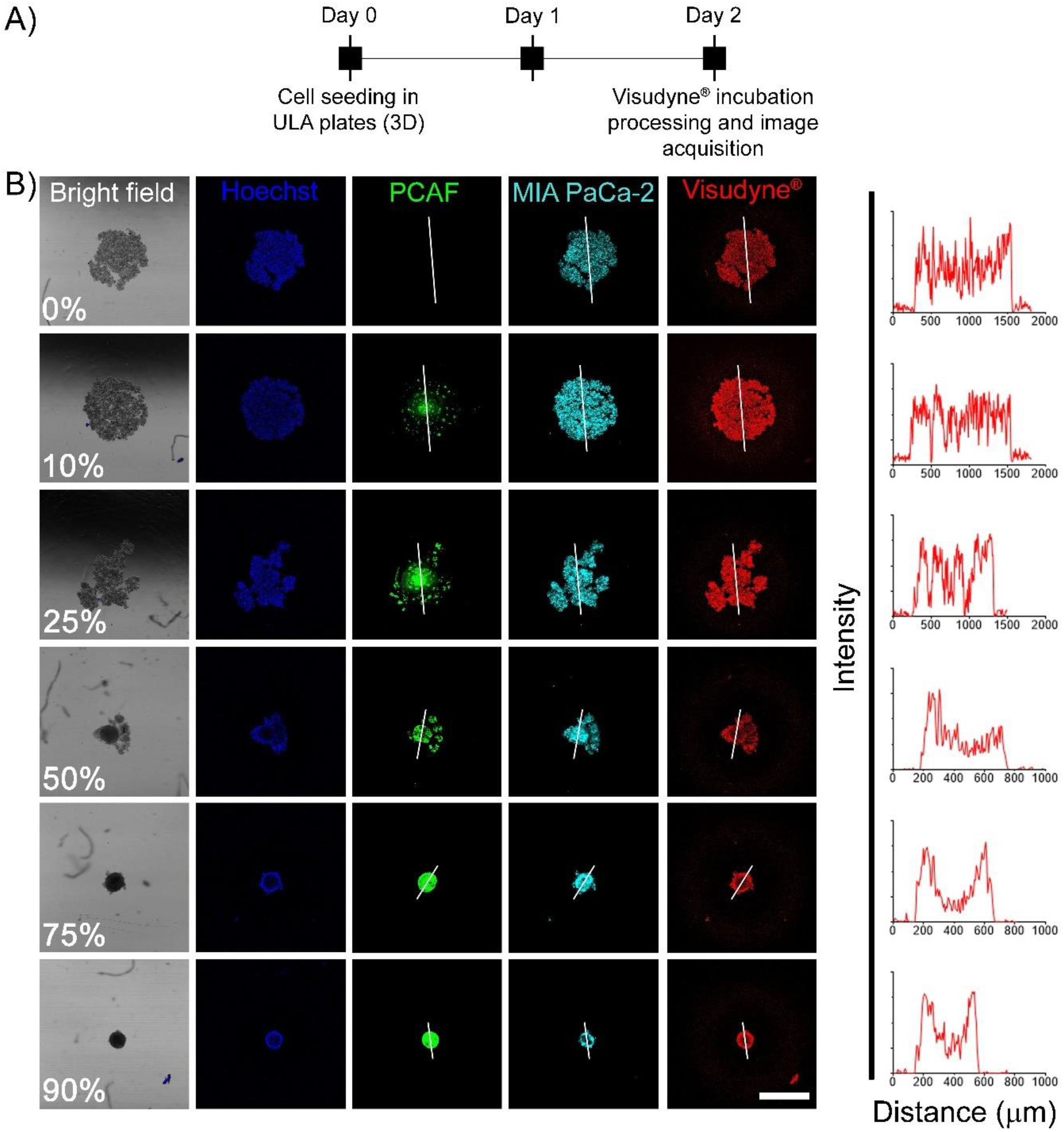
Visudyne® administered photosensitizer distribution in heterotypic 3D spheroids. (**A**) Experimental schedule for studying BPD uptake and distribution in homotypic and heterotypic spheroids using PICs. Cells were seeded in CellCarrier Spheroid ULA 96-well Microplates followed by incubation with Visudyne® for 90 min. The spheroids were then washed, fixed in PFA, counter-stained with Hoechst and visualized under a confocal fluorescence microscope. (**B**) Representative confocal images of the central optical section of the 3D spheroids. Mia PaCa-2 cells and PCAFs are pseudo-colored as cyan and green. Nuclei are stained with Hoechst (pseudo-colored as blue) and BPD is shown in red. Profile plots were generated using Image J to monitor BPD distribution across the central plane. As suggested by the profile-plots, the distribution of BPD was homogeneous for spheroids generated with 0%, 10% and 25% PCAFs. However, as the PCAF percentage increased beyond 50% the distribution of BPD appeared to be heterogeneous with higher signal on the periphery and lower signal in the spheroid core (scale bar = 500 µm).

## Discussion

The desmoplastic response observed in PDAC tumors is a major impediment, which creates mechanical and biological barriers to therapies^51^. Reports suggest that the stromal components (including CAFs and extra-cellular matrix) can account for as much as 90% of tumor volume^11,12^.

Most therapies for PDAC ultimately fail and the 5-year survival rate of 7% has not changed in recent decades^1,52,53^. In this context, PDT offers an attractive alternative due to its ability to modulate the stroma and cancer cells in the PDAC microenvironment^19,23^. PDT has been previously employed in the clinic and has shown to successfully induce necrosis in the irradiated regions of PDAC tumors^28,29^. While these studies were performed using Visudyne®, and although the stromal content of these tumors was not quantified, the results were encouraging. In the present study, we provide a comparison of Visudyne®-PDT and a clinically emergent targeted photosensitizer construct, referred to as PIC on *in vitro* PDAC spheroids with different ratios of Mia PaCa-2 and PCAFs. These spheroids more faithfully recapitulate the inter-patient heterogeneity observed in PDACs and enable longitudinal monitoring of cellular viability in response to PDT. We employed Cetuximab, a clinically approved anti-EGFR antibody, previously utilized for guiding fluorescence resection surgeries in pancreatic and other tumors (NCT03384238)^24,38,40,54,55^. Moreover, PICs for PIT, using the photosensitizer IRDye700 have recently been approved in Japan as Cetuximab sarotalocan for the treatment of head and neck cancers and is currently in Phase III trials in the US (NCT03769506)^34,48,49^. In this study, we demonstrate a higher efficacy of PIT, as compared to Visudyne®-PDT, specifically on tumor spheroids with more than 50% PCAFs. The higher efficacy of PIT could probably be due to a more homogenous distribution of the antibody conjugate in the heterotypic PDAC spheroids, as suggested by confocal fluorescence studies (**Figure 7**). In contrast, an uneven (heterogeneous) distribution of the photosensitizer as delivered through Visudyne® led to the PS deposition in the cells constituting the surface layers of the spheroids, as suggested by Confocal fluorescence microscopy studies (**Figure 8**). Aside from variations in PS penetration, differences in the mode of action of Visudyne®-PDT and PIT may result in the activation of different cytotoxic and/or stress response pathways thereby accounting for the differences in viability observed for the two treatments. PCAFs in general have been reported to confer treatment resistance in PDAC and the relatively lower response of Visudyne®-PDT in the spheroids with higher PCAF percentage may likely be due to the treatment resistance conferred by PCAFs. The fact that the spheroids with higher PCAF percentage show an increase in their relative viability upon Visudyne®-PDT treatment suggests that there is also a mechanistic resistance induced by the high PCAF content, which is not effective at limiting PIT phototoxicity. Although preliminary, these results highlight that PIT is agnostic to desmoplasia-induced resistance and further in vivo preclinical investigation is warranted.

Although PIT was observed to be more effective irrespective of the PCAF percentage, it was also found to be phototoxic to PCAFs (specifically in spheroids with higher PCAF percentage) in the heterotypic spheroids (**Figure 5**). This is likely due to the expression of EGFR on PCAFs, although at a lower level than Mia PaCa-2 cells^23^. It has been long known that stromal components, including CAFs, in PDAC serve a complex role that complicates potential treatment options^56^. Several studies have focused on therapeutic strategies targeting CAFs or the Extracellular Matrix (ECM) to decrease desmoplasia and enhance the efficacy of treatments such as chemotherapy^6,57^. Interestingly, many of these pre-clinical and clinical studies revealed that depletion of stroma can result in more aggressive tumor growth, with one study in particular finding that depletion of stroma and myofibroblasts in a mouse model resulted in increased invasiveness and reduced survival rates^57^. This suggests that the stroma may have both tumor-suppressing and tumor-promoting roles and that by destroying the stroma, critical mechanisms of tumor growth and metastasis are diminished^8,9,58^. Emerging data suggests that the high heterogeneity and plasticity in PCAFs where targeting of specific PCAF sub-populations may provide survival benefits^59-62^. Moreover, EGFR expression has been reported to increase with the activation of CAFs, making them a viable target for EGFR-specific PIT^63^.

## Conclusion

PDACs are notoriously tough to treat, and their 5-year survival is amongst the lowest of all cancer types. The high stromal content with significant intra- and inter-patient heterogeneity has been recognized as a major factor in treatment failures. In this study, we show a high phototoxicity of PIT, on heterotypic PDAC spheroids developed with high PCAF percentage. In contrast, the clinically approved photosensitizer formulation, Visudyne®, was not as effective in spheroids with high PCAF percentage. With several clinical trials on antibody-fluorophore conjugates for guiding tumor resection surgeries and the recent clinical advances of PIT for head and neck cancers, this study puts forward compelling evidence for the efficacy of PICs in PIT of highly desmoplastic pancreatic tumors. Moreover, given the relatively moderate to low success of most treatment options for pancreatic cancers, PIT may provide a novel method for pancreatic cancer treatment and sensitization to combinatorial therapies.

## Acknowledgements

This work was supported by grants from National Institute of Health P01 CA084203, R01 CA231606, R01 CA266855 and R01 CA260340 to TH, and K99CA215301 to GO.

